# Passive plasma membrane transporters play a critical role in perception of carbon availability in yeast

**DOI:** 10.1101/2021.09.11.459425

**Authors:** Amogh Prabhav Jalihal, Christine DeGennaro, Han-Ying Jhuang, Nicoletta Commins, Spencer Hamrick, Michael Springer

**Affiliations:** Department of Systems Biology, Harvard Medical School

## Abstract

Recently, our lab found that the canonical glucose/galactose regulation pathway in yeast makes the decision to metabolize galactose based on the ratio of glucose to galactose concentrations in the external medium. This led to the question of where and how the ratio-sensing is achieved. Here, we consider the possibilities of an intracellular, extracellular, or membrane bound ratio sensing mechanisms. We show that hexose transporters in the plasma membrane are mainly responsible for glucose/galactose ratio-sensing in yeast. Further, while the glucose sensors Gpr1, Snf3, and Rgt2 are not required for ratio sensing, they help modulate the ratio sensing phenotype by regulating the expression of individual transporters in different environments. Our study provides an example of an unexpected, but potentially widespread, mechanism for making essential decisions.

## Introduction

Organisms respond to their environments by integrating multiple signals relevant to their metabolism [1][2] and growth[3] through regulatory networks. In the natural environment, cells often have to make decisions about which nutrients to utilize, and therefore which metabolic genes to express. The galactose metabolic genes (GAL genes) in yeast are a canonical example; until recently, it was believed that these genes are only expressed if the concentration of a preferred sugar, glucose, in the environment falls below a certain threshold. We recently discovered that the situation is more complex than this, and that the decision to induce these genes is in fact regulated by the ratio of the concentrations of galactose and glucose [4]. Although this result is surprising in the context of previous beliefs about catabolite repression, it is less surprising when one considers the potential fitness effects of choosing to use a less preferred sugar when it is highly abundant. In this case, the negative fitness effects of using cellular resources to express the GAL genes when they are not absolutely essential would be balanced against the positive fitness effects of having access to a large additional pool of nutrients [5]. Indeed, we were able to show that ratio sensing can confer a fitness advantage to yeast grown in a mixture of galactose and glucose [4].

Ratio sensing may be advantageous in many other contexts, and has been observed in a few biologically important settings. Widely known examples include balance among adenine nucleotides regulating metabolism [6], the X-to-autosome ratio determining sex in *Caenorhabditis elegans* [7], and the bone morphogenetic protein signaling pathway in mammalian cells responding to ratios of ligands [8]. All of these examples, due to the crucial roles they play, are difficult contexts in which to disentangle the mechanism by which ratio sensing is achieved. Model systems such as the GAL pathway in yeast by virtue of being non essential and genetically tractable promise to offer insights into the type of mechanisms that can give rise to ratio sensing.

We set out to investigate the mechanism of ratio sensing in the GAL pathway. Our previous work identified the location of the sensing mechanism as being upstream of the canonical GAL signaling pathway [4]. We systematically separate between potential mechanisms by which ratio sensing could occur, i.e. inside the cell, outside the cell, and at the cell membrane via the hexose transporters. We demonstrate that hexose transporters are the site of galactose/glucose ratio sensing. Finally, we investigate the role of glucose sensors in fine tuning the ratio sensing phenotype in response to various carbon stress conditions. We argue that nutrient transporters have an important and overlooked signal processing role in the control of sugar metabolism, and possibly in many other contexts.

## Results

We previously established a platform for assessing galactose utilization in yeast ([4], Figure 1A, B). Briefly, a YFP gene is placed under the control of the *GAL1* promoter on the *HO* locus. Galactose induces the expression of *GAL1* through Gal4, and glucose represses its expression level via Mig1 (Figure 1C). The GAL pathway induction can be quantitatively assessed by flow cytometry. As the GAL pathway can respond in a bimodal manner across a population [4], we monitor both the fraction of cells that are induced and the mean induction level of the induced cells. We can thus generate a matrix of the induction profile of a strain in various mixtures of glucose and galactose (Figure 1 B). Using this platform, we aim to understand the point(s) where ratio-sensing occurs. Ratio-sensing can happen whenever there is competitive binding of substrates or proteins on the same sites. Alternatively, there could also be a dedicated mechanism for the ratio-sensing. We reasoned that the galactose/glucose ratio-sensing could be achieved (a) intracellularly via competitive binding of sugars to certain proteins, such as Gal3, or (b) extracellularly via specialized sugar sensors, or (c) at the cell membrane via competitive binding of sugars to hexose transporters (Figure 1D). In the following sections we systematically investigate these three possible possible mechanisms of galactose/glucose ratio sensing.

**Figure 1:**
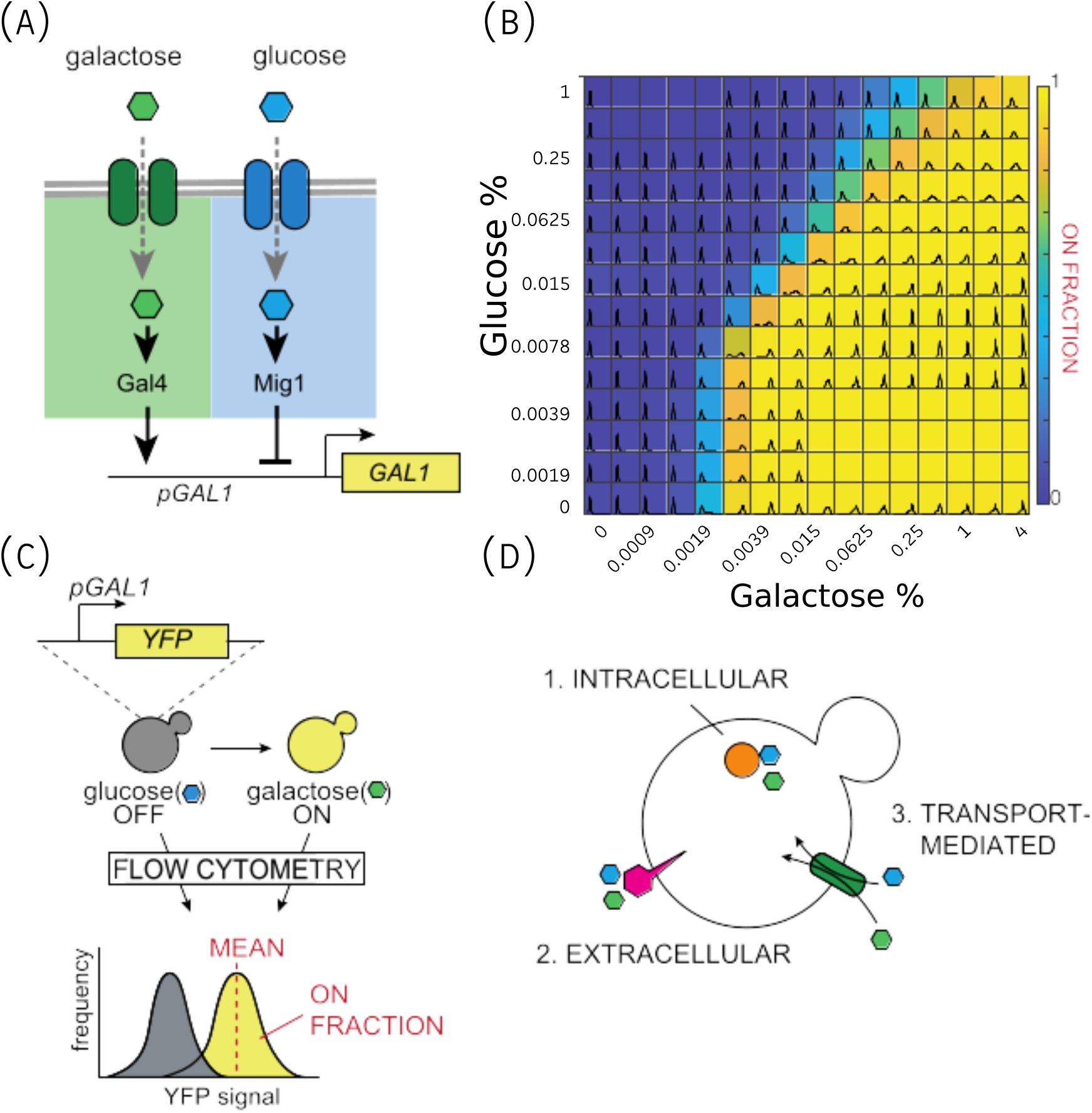
GAL metabolic genes respond to a ratio of glucose and galactose. (A) Schematic of experimental setup. Strains expressing YFP under the *GAL1* promoter are used as the read out GAL pathway activity. When cultured in glucose media, *GAL1p-YFP* is not expressed; conversely, when cultured in galactose media, *GAL1p-YFP* is expressed at a high level. YFP levels were measured by flow cytometry. (B) Heatmap of the fraction of cells expressing *GAL1* above basal levels in a galactose/glucose double-gradient setting [4]. The gradient is constructed as a two-fold dilution series starting from 4% wt/vol galactose in the column on extreme right and 1.0% wt/vol glucose in the top row. The left extreme column and bottom row contain 0% galactose and glucose respectively. Labels on the axes indicate concentration of the sugar in percent wt/vol. (C) Simplified diagram of the GAL pathway. Galactose activates the *GAL* pathway though Gal4, while glucose represses the *GAL* pathway though Mig1. (D) Three possible ratio-sensing regimes. (1) ratio-sensing is achieved via an unknown intracellular mechanism, (2) extracellular sugar sensors, or (3) competitive transport through shared transporters.

### Internal glucose levels have little effect on galactose/glucose ratio-sensing

Both intracellular and extracellular glucose have the potential to affect the GAL pathway induction. Distinguising between whether the ratio of glucose and galactose is sensed internally, at the plasma membrane, or externally is difficult as changing the external glucose or galactose concentrations changes the flux through the membrane and the internal concentrations of both glucose and galactose. To circumvent this problem we used maltose instead of glucose. After uptake, the disaccharide maltose is hydrolysed into two glucose monosaccharides ([9] and Figure 2A). Furthermore, maltose is imported through specific transmembrane transporters for di-saccharides and not through hexose transporters [10] [11]. Thus maltose could be used to distinguish intracellular versus extracellular galactose/glucose ratio-sensing. We measured Gal1pr-YFP induction in yeast cells grown in a galactose/maltose double-gradient. In contrast to a glucose/galactose double gradient, the GAL pathway induction responded solely to the galactose levels and not the galactose/maltose ratios. The lack of inhibition of the GAL pathway by maltose was not due to insufficient internal glucose levels. The level of maltose added was sufficient to raise internal glucose levels to the point that Mig1 was localized to the nucleus [12] (Figure 2B-C). While clearly distinguishable from the inhibitory role of glucose, at high levels of maltose and levels of galactose close to the induction threshold with no competing sugar, maltose had a small inhibitory role of Gal1pr-YFP induction (Figure A.1). Based on these results we ruled out internal glucose levels as the dominant factor in galactose/glucose ratio sensing.

**Figure 2:**
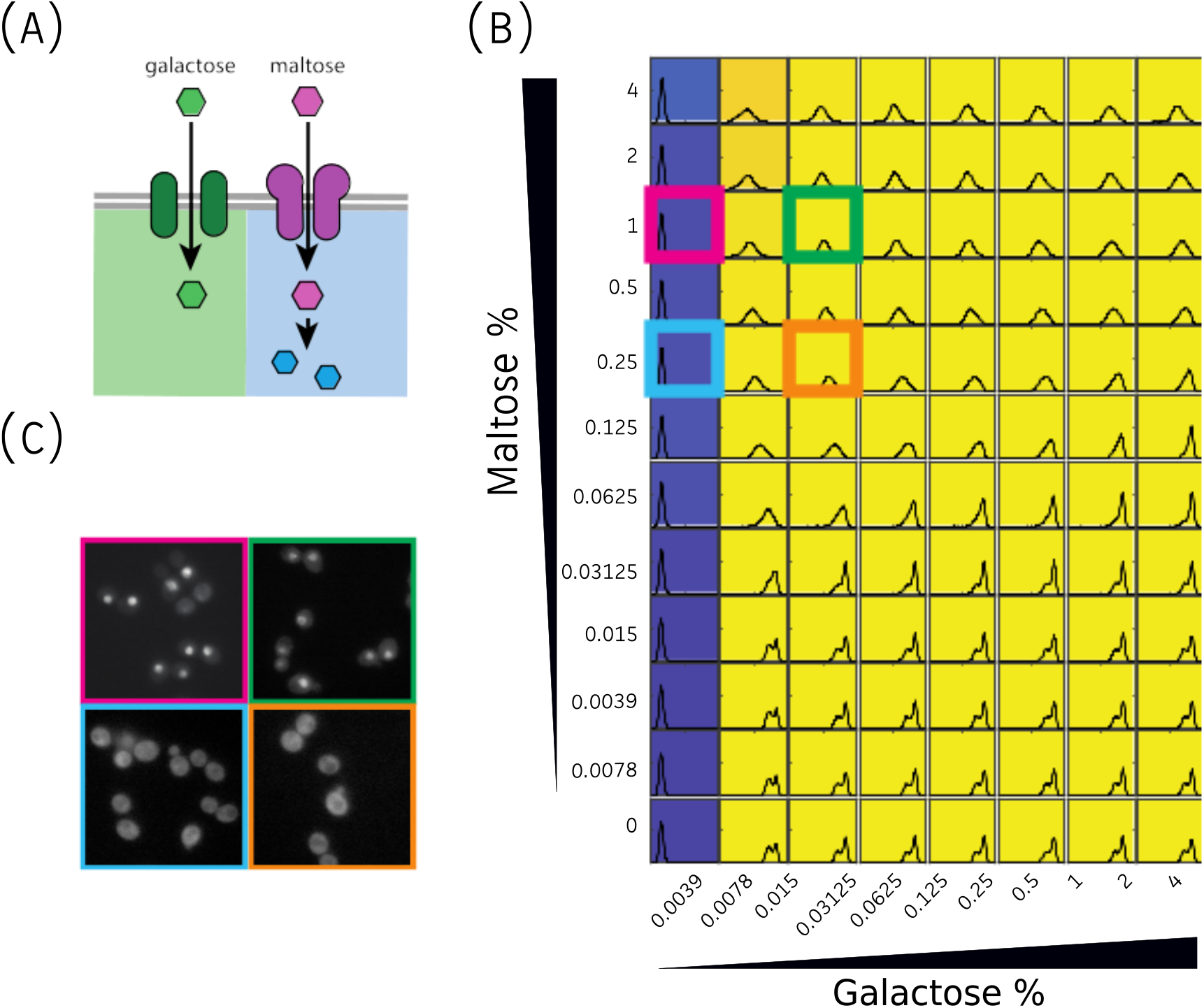
Intracellular glucose signaling is not sufficient for the ratio-sensing functionality. (A) To test whether the ratio-sensing happens intracellularly, we took advantage of the sugar maltose, which is transported independently from galactose and is hydrolyzed into two units of glucose inside the cell. (B) Ratio-sensing is not evident in a galactose/maltose double gradient. Labels indicate the concentrations of sugars in % wt/vol (C) Localization of Mig1-YFP fusion protein in the given sugar conditions marked in (B). Nuclear localization of Mig1 (in 1% maltose) indicates glucose-mediated repression of *GAL* and hence the conversion of intracellular maltose to glucose.

### Plasma membrane glucose sensors are not required for galactose/glucose ratio-sensing

As intracellular glucose has little effect on the galactose/glucose ratio-sensing behavior, we next asked if sugar sensors on the plasma membrane act as the source of the ratio-sensing behavior. There are believed to be three glucose sensors on the yeast plasma membrane — Snf3, Rgt2 and Gpr1 ([13] and Figure 3A), and no known plasma membrane galactose sensors. Snf3 and Rgt2 are known to regulate glucose transporter levels [14]. Deletion of these sensors would also change transporter levels and thus make interpretation difficult. In order to remove the confounding regulatory effects of the sensors on the hexose transporter levels, we used a strain in which all the hexose transporters had been deleted [15] where a single hexose transporter (( HXT2)) had been reintroduced under constitutive expression. Deletion of the three glucose sensors (‘triple-sensorless’) in this single hexose transporter background had a minimal effect on the ratio-sensing decision front (Figure 3B-C), arguing that these glucose sensors are not the source of the galactose/glucose ratio-sensing behavior. Notably, a more pronounced GAL-off population can be seen in mixtures of high glucose and galactose in the strain with the sensors, suggesting a role of glucose sensors on the fraction of induced cells (Figure 3D). Nevertheless, the effect of the deletion of glucose sensors on the decision front is minimal.

**Figure 3:**
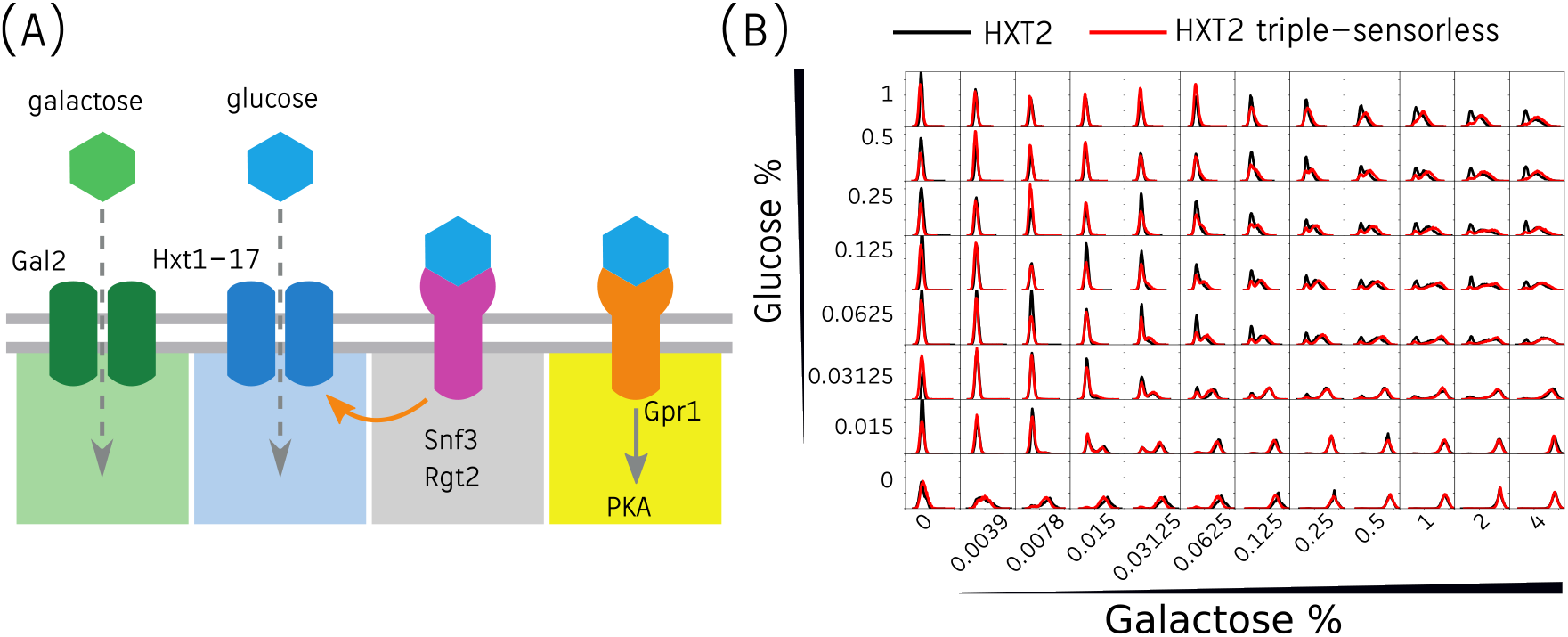
Extracellular glucose sensors are not required for the ratio-sensing behavior. (A) Three known glucose sensors -- Snf3, Rgt2, and Gpr1, sense extracellular glucose concentration. Snf3 and Rgt2 are known to regulate the expression of hexose transporters. Gpr1 regulates the PKA activity. (B) Overlay of induction profiles of the single-HXT2 strain (black) and *snf3*Δ *rgt2*Δ *gpr1*Δ triple-sensorless single-HXT2 strain (red).

### Plasma membrane transporters affect galactose/glucose ratio-sensing

We have thus far ruled out intracellular mechanisms and plasma membrane glucose sensors as the main source of the galactose/glucose ratio-sensing behavior. This strongly implicates plasma membrane transport as the critical mechanism underlying ratio-sensing. Ratiometric response would be achieved by competition between galactose and glucose for transport by hexose transporters (Figure 4A).

**Figure 4:**
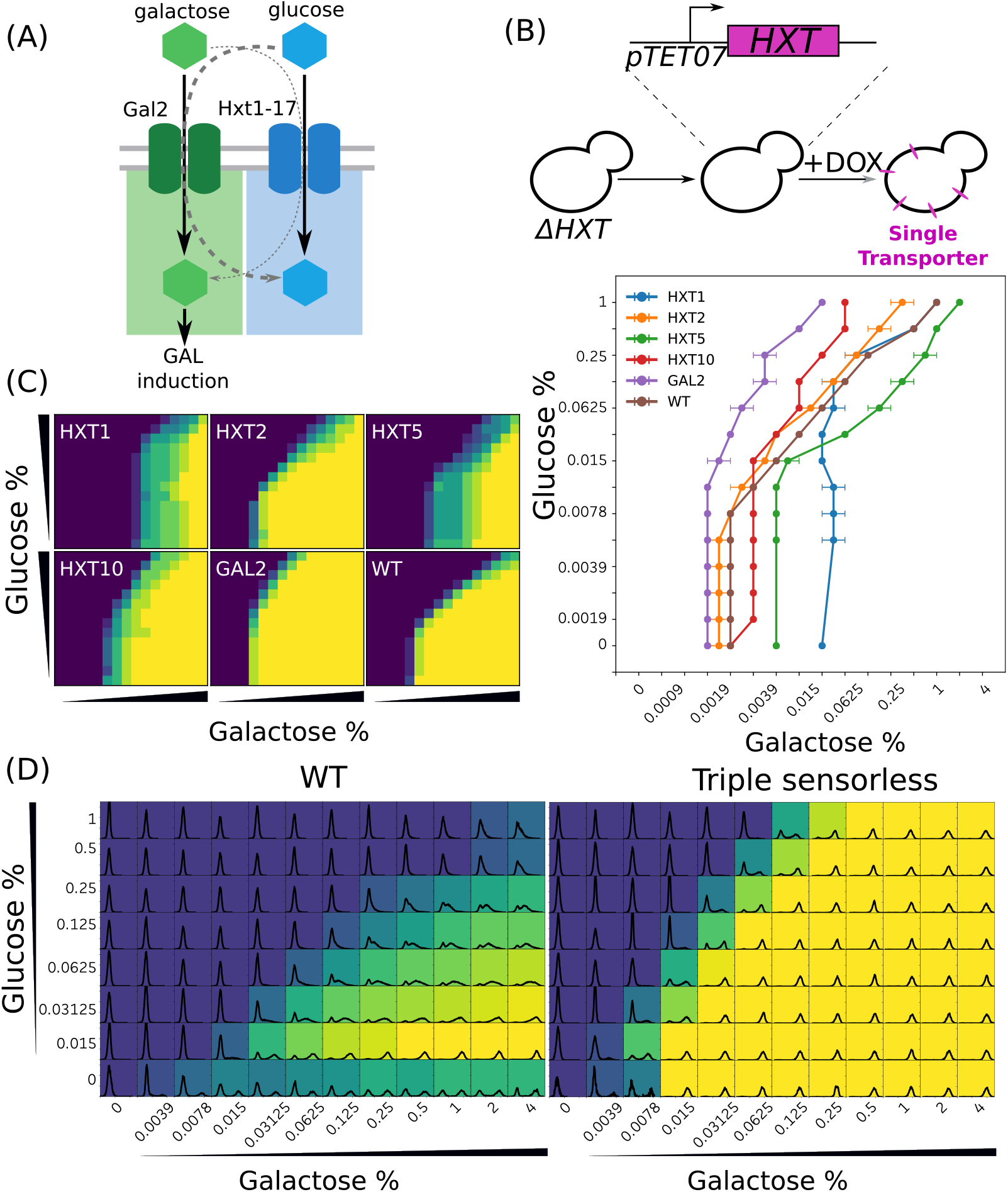
Properties of hexose transporters determines the ratio-sensing behavior. (A) Hexose transporters have differential affinities for galactose and glucose. (B) Single hexose transporters were individually expressed under the *TETO7* promoter in a hxt-null strain where *HXT1-17* and *GAL2* were knocked-out. Doxycycline was used to induce the expression of single hexose transporters before conducting galactose/glucose double-gradient experiments. (C) Heatmaps (left) and thresholds (right) of the GAL induction profiles of each single-transporter strain. Gradients were constructed as decribed in Figure 1. (D) Heatmaps showing the effect of the hexose sensors on the decision threshold. Histograms show the WT response on the left and the triple-sensorless strain on the right.

To test this hypothesis, we generated several single transporter strains; strains in which all hexose transporters were knocked out (*HXT1-17* and *GAL2*), and one of the hexose transporters was introduced back under a constitutive promoter (Figure 4B). If hexose transporters are the source of the ratio-sensing behavior, the difference in their affinity to different sugars should give rise to different ratio-sensing thresholds – i.e. the ratio of the two sugars where half of the cell population activates the GAL pathway. Remarkably, we observe significant differences among the induction profiles of different single hexose transporter strains, suggesting that the plasma membrane hexose transporters as the main source of the ratio-sensing behavior, consistent with the hypothesis of competitive transport (Figure 4C).

Plasma membrane glucose sensors are known to regulate hexose transporter levels [14]. While our results showed that plasma membrane glucose sensors are not required for the galactose/glucose ratio-sensing behavior, given their role in regulating transporter levels, we should expect deletion of the glucose sensors in a strain not deleted for hexose transporters to have an altered ratiometric response. As expected, the deletion of glucose sensors shifted the ratiometric response (Figure 4D).

### Ratio sensing phenotype and hexose transporter expression are strongly correlated to carbon conditions

In the previous section we provided evidence that the plasma membrane hexose transporters can affect glucose-galactose ratio sensing. Furthermore, the observations from the triple-sensorless strain suggested that the cells could regulate the characteristics of the ‘decision boundary’, i.e. the ratio at which the GAL genes are turned on, by changing transporter expression in different conditions. We tested two predictions that would occur if hexose transporter levels were differentially regulated and hexose transporters were responsible for ratiometric sensing - First, changing the effective *K*_*m*_ of the transporters should affect the gal ratiometric response. Second, the cell’s perception of the current carbon sufficiency should affect membrane transporter repertoire. In order to study the first possibility, we measured the decision boundary and the hexose transporter expression in a variety of rich (2% glucose, 0.2% glucose, 2% fructose, 2% maltose) and poor (2% raffinose, 0.02% glucose, 2% acetate) carbon sources. The rationale is that cells should modulate their transporter composition based on the carbon composition of the environment, which should lead to changes in the effective *K*_*m*_ for glucose and galactose. This change in *K*_*m*_ would alter the position of the decision front. We carried out these measurements in three strains, a triple-sensorless strain, a wildtype strain with a CEN.PK-2 background, and a wildtype strain with the S288c background. We observe that in the wildtype strains, the decision boundaries depend on the quality of the carbon source, whereas the triple-sensorless strain shows no variation in the decision boundaries (Figures 5 (A), (B), (C)). Further, performing a Principal Component Analysis (PCA) on the population histograms from all the glucose-galactose ratios measured, we observe that the wildtype strains are separated based on the quality of the carbon source, whereas the sensorless strain displays no such variation (Figure 5 (D)). By examining the *HXT* expression (Figures 5 (E), (F), (G)), and performing a PCA on the transcript counts of the 16 *HXT* genes (Figure 5 (H)), we find a similar separation of conditions by carbon quality in the wildtype strains which is absent in the sensorless strain. Finally, we observe that the clusters produced by the phenotypic data are highly correlated with those produced by the transcriptomic data (Figure 5 (I)). Together, this data offers support for our hypothesis that the *GAL* response is modulated in response to external carbon sources via the regulation of plasma membrane transporter composition.

**Figure 5:**
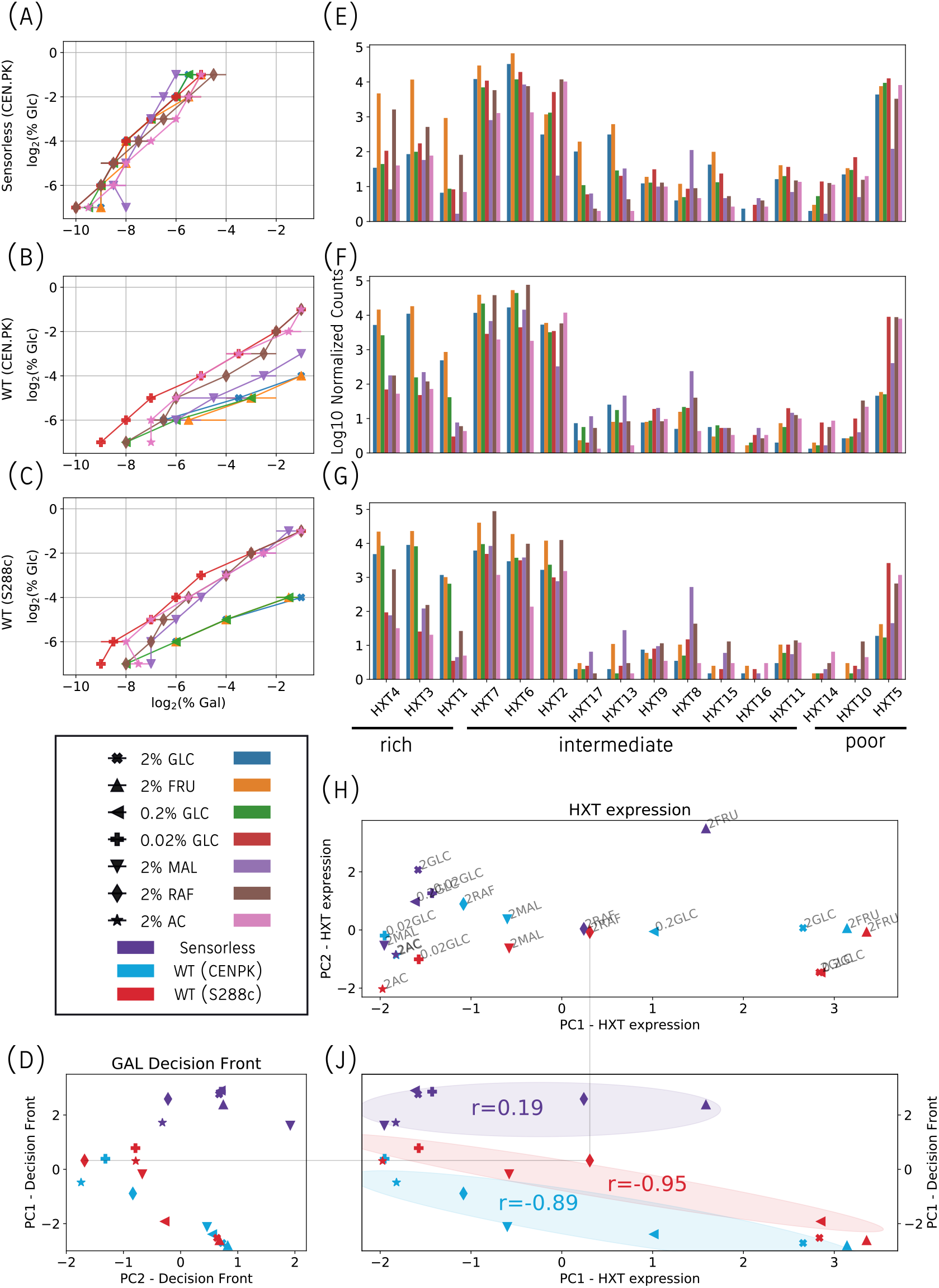
Comparing the decision front phenotype and HXT expression across carbon conditions. Colors and shapes used to distinguish carbon conditions and strains are explained in the legend. (A), (B), (C) Decision boundaries of the triple-sensorless, WT CEN.PK-2, and WT S288c strains grown in a seven different carbon conditions. (D) PCA plot of the phenotypic data. The PC1 axis (the principal component capturing the highest variation in the data) is represented along the *y*-axis. Colors represent the three strains. (E), (F), (G) Barplots showing the log10 of normalized counts of 16 hexose transporters across the carbon conditions studied. The HXTs along the *x*-axis are arranged according to their loading in the first principal component in (H). (H) PCA plot of HXT expression. The PC1 axis falls on the *x*-axis. (I) Plot comparing the PC1 coordinates from the (D) on the *y*-axis to those of (H) on the *x*-axis. Highlighted region shows the spread of points of each strain. The gray line indicates the comparison of one example condition, 2% Raffinose in the S288c background. Overlayed text shows the Pearson correlation values of two PC1 coordinates. Note that the coordinates of the two wildtype strains have significant correlations (p ¡ 0.05), while the correlation of the sensorless strain is not statistically significant.

We investigate the second prediction by considering the effect of maltose on our three strains. The S288c background which is defective in maltose uptake [16] would be expected to exhibit a response similar to other poor carbon conditions, while the CEN.PK-2 background which can import maltose should exhibit a phenotype similar to growth in rich medium. In fact, both of these predictions are borne out from the data, as seen in Figure 5(D). Importantly, we observed that the composition of HXT transcripts in both these strain backgrounds is very similar (5(H)), indicating that the sensors regulate HXT gene expression, while other factors (like growth rate) might regulate HXT protein levels.

## Discussion

We have previously shown that ratio sensing is achieved upstream of the canonical GAL pathway [4]. In this study, we have provided evidence that the hexose transporters are the site of ratio sensing. Neither changes in intracellular glucose level, nor deletion of plasma membrane glucose sensors affect the galactose/glucose ratio. Instead, changing the composition of plasma membrane transporters changes the ratiometric decision front whether done directly through synthetic over-expression or through native regulation in different carbon sources.

Previous work has shown that cells optimize global translation under various growth conditions by modulating tRNA pools to match codon usage patterns [17]. Our observation that the ratio sensing phenotypes and transporter expression can be clustered into ‘high’ and ‘low’ energy states can potentially indicate another manifestation of this global cellular state. Cells might ‘lock in’ to hexose transporter expression patterns that might be optimal to support global cellular energy needs.

While we did not directly measure the effective *K*_*m*_ values of the *HXT* s for galactose, our hypothesis is that differences in the effective *K*_*m*_ account for the differential effects of the HXTs on the decision fronts. A recent study by Montano-Gutierrez *et al.* has demonstrated the complex regulation of the HXTs by various glucose dependent transcription factors using the framework of rate-affinity tradeoff[18]. Montano-Gutierrez *et al.* propose that the HXTs might be regulated according to their capacity to maintain optimal glucose flux in response to varying environmental glucose levels. Our study of glucose-galactose ratio sensing, particularly the observation that HXT expression is correlated with ratio sensing phenotype, shows that there are considerations beyond optimal glucose transport in the regulation of HXT expression.

Transporters have received a lot of attention in recent years, from the so-called ‘dual-transporter’ systems of phosphate and zinc in yeast [19], to the recent characterizations of the rate-affinity tradeoffs that shape transporter expression [20]. Our work showing competitive transport as a mechanism of ratio-sensing expands this literature. In the context of yeast, other nutrient transporter systems such as the amino acid transporters are a possible target system in which to explore transporter-based ratio sensing [21]. This paper will hopefully serve as a guide for exploring rationmetric sensins in other pathways and systems.

## Methods

### Growth Conditions and Media

Cells were grown for 16–18 h in synthetic minimal media with 2% (wt/vol) raffinose, to an OD of ~0.1, and then diluted 1:1000 in mixtures of sugars. Cells were grown for 8 h at 30°C in flasks or 96-well plates and then washed twice in TE (10 mM Tris, 1 mM EDTA, pH 7.5) in preparation for flow cytometry. Cells to be imaged by microscopy were transferred to a micro-well plate (Matriplate from Metrical Bioscience) coated with concanavin A grown in synthetic media with different mixtures of glucose and galactose.

### Flow Cytometry

Flow cytometry was performed on a Stratedigm S100EX with A700 automated plate handling system.

### Fluorescence Microscopy

Images were captured with a Ti Eclipse inverted Nikon microscope using Micromanager. A Hamamatsu Orca-R2 camera was used to capture fluorescent images and a Scion CFW-1612M for bright-field images. Nuclear and cytoplasmic of Mig1p-GFP localization was analyzed using custom MATLAB software.

### Data Analysis and availability

Analysis of flow cytometry data was performed using custom-written MATLAB code (available upon request). A 2D Gaussian mixture model fit to the YFP populations was used to identify mean expression (‘on’ level) and the fraction of population expression YFP (‘on’ fraction). Processed data files and scripts to generate figures are available here. In order to carry out PCA on the phenotypic data (Figure 5 (D)), the population histograms for the 96 well plates were first processed in order to obtain the ‘on’ fraction metric. The vector of 96 values for each strain and growth condition was used as the feature vector while performing the PCA.

### RNA sequencing

#### RNA extraction and library preparation

Cells were grown overnight in YPD, washed twice with minimal media, and then inoculated into the target media (2% glucose, 0.2% glucose, 0.02% glucose, 2% raffinose, 2% acetate, 2% fructose, 2% maltose, or 2% galactose). These cultures were allowed to grow up to an OD of 0.5 or up to 24 hours in case of slow growing cultures. The cultures were then spun down and the pellets were snap-frozen with liquid nitrogen and store at −80°C. Total RNA was then extracted using acid phenol-chloroform extraction as follows: frozen pellets were resuspended in equal amounts of TES buffer (10 mM Tris-Cl pH 7.5, 10 mM EDTA, 1% SDS) and pre-heated acid phenol:chloroform:IAA (125:124:1, ThermoFisher Scientific, Catalog Number: AM9722), and were incubated at 65°C for 30 min with shaking. The supernatants were cleaned with acid phenol:chloroform:IAA again and RNA was precipitated with 0.3M NaOAc and 75% EtOH at pH 5.5 at −80°C overnight. Samples were cleaned with 80% cold EtOH and resuspended in RNase-free water, treated with DNase I at 37°C for 30 min and finally cleaned with the miniElute kit (QIAGEN, Catalog Number: &28004). For next-generation sequencing, the Illumina TruSeq Stranded mRNA Library Prep kit was used (Catalog Number: &20020594). Library preparation was performed according to the instructions provided with the kit. The TruSeq RNA adapter plate, 96 plex plate was used for indexing (Part Number: &15016427). Sequencing was performed using the NextSeq 500 System.

#### RNA-seq analysis

Raw reads were quantified using Salmon [22]. Data processing was carried out using DeSeq2 [23]. Principal component analysis was carried out using Scikit-Learn Python library [24].

## Acknowledgements

APJ would like to acknowledge Sarah Boswell for her help with library preparation. This work was supported by NIH Grant R01 GM120122-01 and NSF Grant 1349248.

## A Supplementary material

**Table 1:** Strains used in this study. All strains with single HXTs and sensorless strains are derived from the CENPK strain background.

| Name | ID | Description |
| --- | --- | --- |
| TetO7-HXT1 | SLYCD109 | MATa leu2-3,112:pTET07:HXT1:LEU2 his3-Δ1 MAL2-8c SUC2 hxt1-17Δ agt1Δ stl1Δ gal2Δ mph2Δ mph3Δ HIS5::PMYO2:rtTA HOΔ::TDH3pr-mCherry:GAL1pr-YFP:NatMX |
| TETO7-HXT2 | SLYCD111 | MATa leu2-3,112:pTET07:HXT2:LEU2 his3-Δ1 MAL2-8c SUC2 hxt1-17Δ agt1Δ stl1Δ gal2Δ mph2Δ mph3Δ HIS5::PMYO2:rtTA HOΔ::TDH3pr-mCherry:GAL1pr-YFP:NatMX |
| TETO7-HXT5 | SLYCD121 | MATa leu2-3,112:pTET07:HXT5:LEU2 his3-Δ1 MAL2-8c SUC2 hxt1-17Δ agt1Δ stl1Δ gal2Δ mph2Δ mph3Δ HIS5::PMYO2:rtTA HOΔ::TDH3pr-mCherry:GAL1pr-YFP:NatMX |
| TETO7-HXT10 | SLYCD117 | MATa leu2-3,112:pTET07:HXT10:LEU2 his3-Δ1 MAL2-8c SUC2 hxt1-17Δ agt1Δ stl1Δ gal2Δ mph2Δ mph3Δ HIS5::PMYO2:rtTA HOΔ::TDH3pr-mCherry:GAL1pr-YFP:NatMX |
| TETO7-GAL2 | SLYCD119 | MATa leu2-3,112:pTET07:GAL2:LEU2 his3-Δ1 MAL2-8c SUC2 hxt1-17Δ agt1Δ stl1Δ gal2Δ mph2Δ mph3Δ HIS5::PMYO2:rtTA HOΔ::TDH3pr-mCherry:GAL1pr-YFP:NatMX |
| WT prGAL-YFP | SLYCD103 | MATa MAL2-8c SUC2 hxt17Δ HOΔ::GAL1pr-YFP:hphNT |
| Triple-sensorless | SLYM01 | MATa MAL2-8c SUC2 hxt17Δ HOΔ::GAL1pr-YFP:hphNT SNF3::natNT2 rgt2Δ gpr1Δ |
| TETO7-HXT2<br>Triple-sensorless | SLYCD123 | MATa leu2-3,112:pTET07:HXT2:LEU2 his3-Δ1 MAL2-8c SUC2 hxt1-17Δ agt1Δ stl1Δ gal2Δ mph2Δ mph3Δ rgt2Δ snf3Δ HIS5::PMYO2:rtTA HOΔ::TDH3pr-mCherry:GAL1pr-YFP:hphNT |

**Figure A.1:**
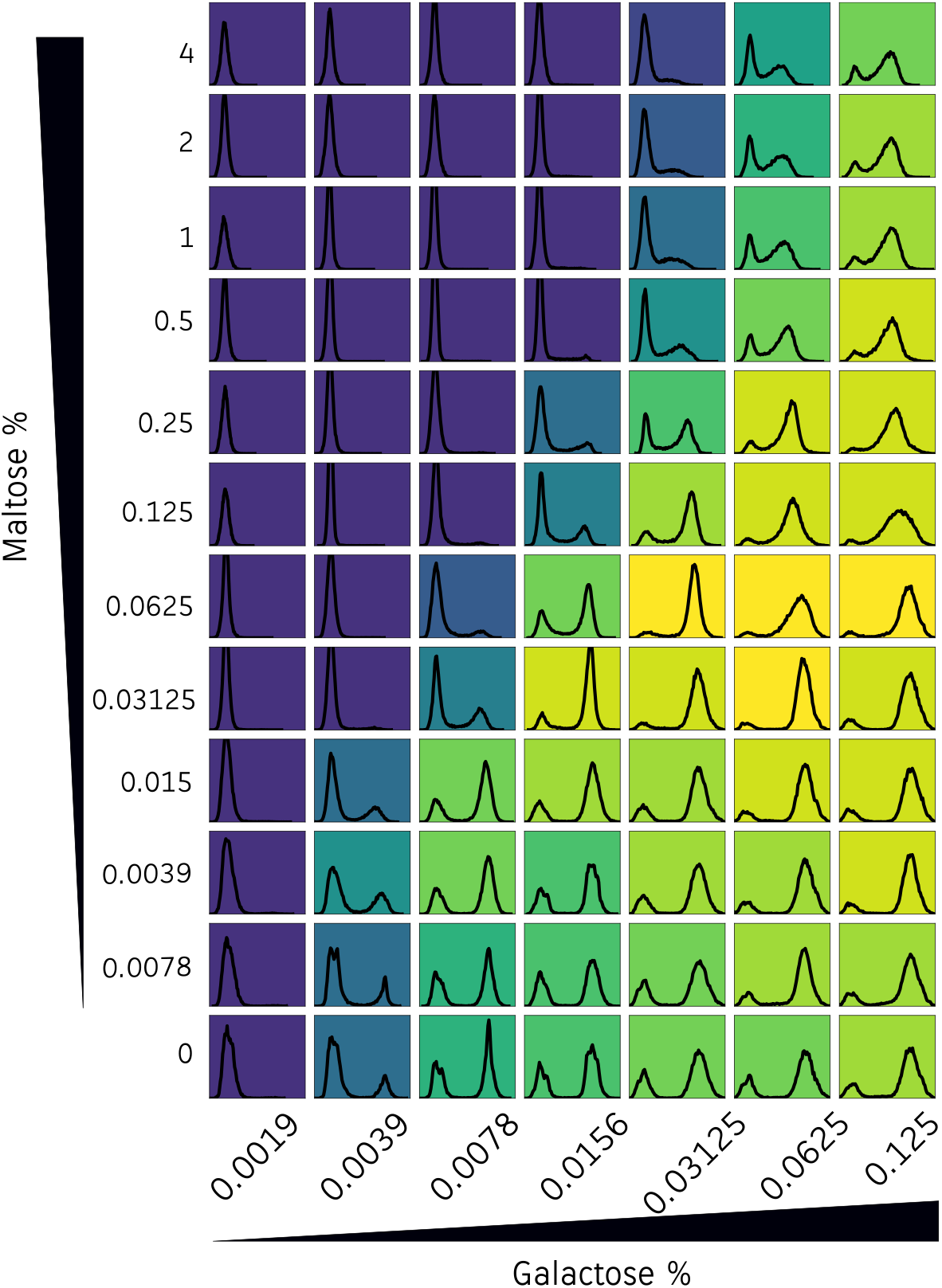
Intracellular glucose signaling is not sufficient for the ratio-sensing functionality. A maltose-galactose double gradient experiment similar to Figure 2B, with a 32X fold lower concentration of galactose (0.125% wt/vol) in the right-most column. We observe no GAL induction below 7.8125e-3%. Repression of GAL induction when galactose concentration is low (< 0.0625%) and the maltose concentration is at least 10X higher than galactose can be observed. However, the behavior is very different from ratio-sensing as the slope of the decision front is much less than 1.

**Figure A.2:**
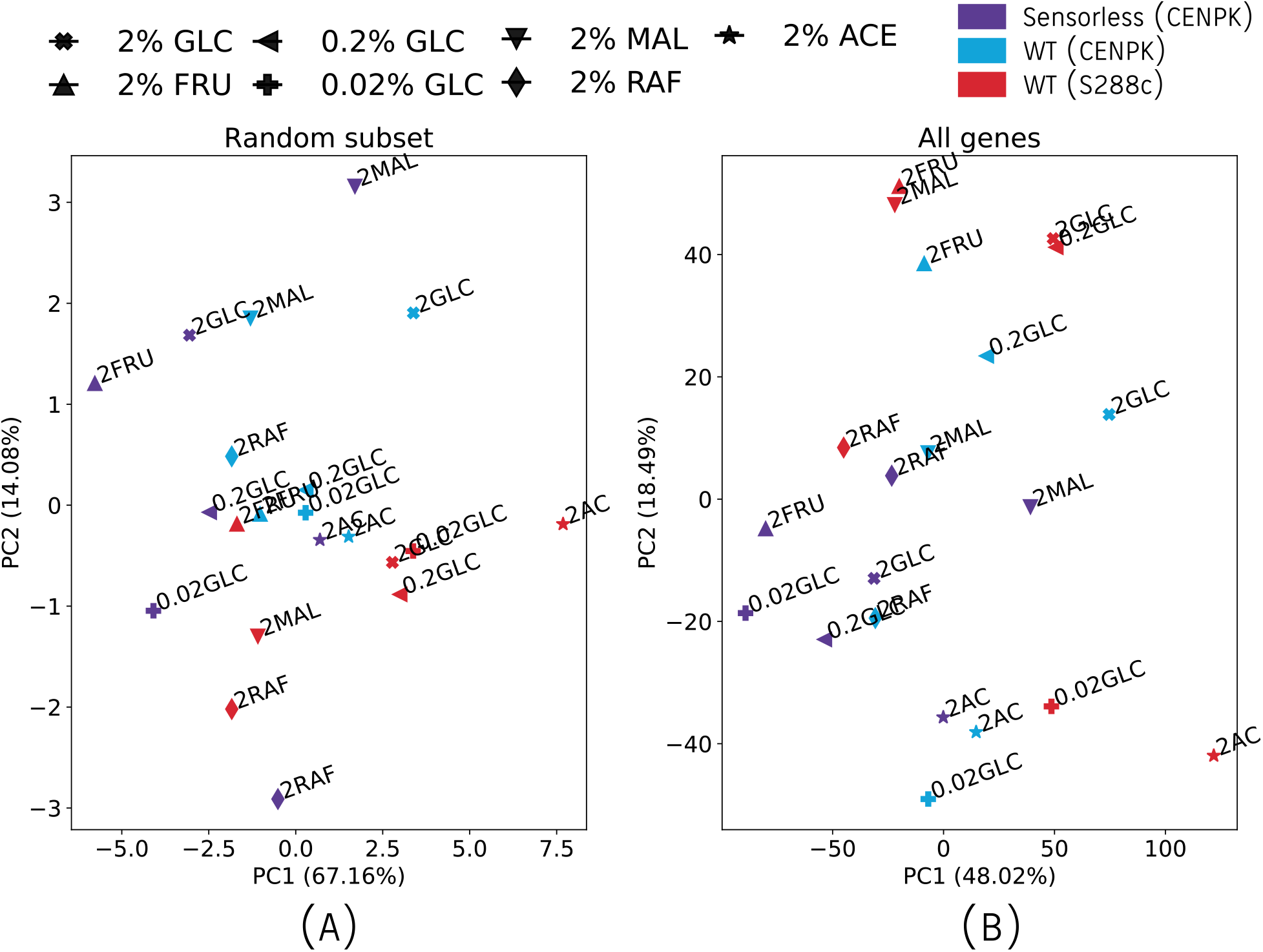
Different choice of genes to compute PCA fails to correlate with carbon condition. The percent variance explained is indicated against the axis labels. (A) Representative random sample of 16 genes (DOG2, PIF1, HTZ1, YOL159C-A, PUT2, GIP1, NOC3, VEL1, SGS1, GLO2, CRG1, SPT16, PNO1, SPO73, PIH1, PHS1). Using this sample of genes, we correlated the Pearson’s correlation value *r* between the PC1 coordinates of nutrient conditions in this plot and those of Figure 5(E), the PCA of the phenotypic data. The correlations are as follows: Sensorless CENPK *r* = 0.55, *p* = 0.2, WT CENPK *r* = −0.31, *p* = 0.49, WT S288c *r* = 0.21, *p* = 0.65. (B) PCA using all 6062 genes as features. The correlation values are: Sensorless CENPK *r* = 0.27, *p* = 0.55, WT CENPK *r* = −0.56, *p* = 0.19, WT S288c *r* = 0.09, *p* = 0.84. We note that for the choice of all genes as features, it appears that the variation in carbon condition is captures along PC2.

## References

[1] Ling Cai and Benjamin P. Tu. Driving the cell cycle through metabolism. Annual Review of Cell and Developmental Biology, 28(1):59–87, 2012.

[2] Victor Chubukov, Luca Gerosa, Karl Kochanowski, and Uwe Sauer. Coordination of microbial metabolism. Nature Reviews Microbiology, 12(5):327–340, 2014.

[3] James R Broach. Nutritional control of growth and development in yeast. Genetics, 192(1):73–105, 2012.

[4] Renan Escalante-Chong, Yonatan Savir, Sean M. Carroll, John B. Ingraham, Jue Wang, Christopher J. Marx, and Michael Springer. Galactose metabolic genes in yeast respond to a ratio of galactose and glucose. Proceedings of the National Academy of Sciences, 112(5):1636–1641, 2015.

[5] Xin Wang, Kang Xia, Xiaojing Yang, and Chao Tang. Growth strategy of microbes on mixed carbon sources. Nature Communications, 10(1):1279, 2019.

[6] Daniel E. Atkinson. Energy charge of the adenylate pool as a regulatory parameter. interaction with feedback modifiers. Biochemistry, 7(11):4030–4034, 1968.

[7] James E Madl and Robert K Herman. Polyploids and sex determination in caenorhabditis elegans. Genetics, 93(2):393–402, 1979.

[8] Yaron E. Antebi, James M. Linton, Heidi Klumpe, Bogdan Bintu, Mengsha Gong, Christina Su, Reed McCardell, and Michael B. Elowitz. Combinatorial signal perception in the bmp pathway. Cell, 170(6):1184–1196.e24, 2017.

[9] R A Dubin, R B Needleman, D Gossett, and C A Michels. Identification of the structural gene encoding maltase within the mal6 locus of saccharomyces carlsbergensis. Journal of Bacteriology, 164(2):605–610, 1985.

[10] Q Cheng and C A Michels. Mal11 and mal61 encode the inducible high-affinity maltose transporter of saccharomyces cerevisiae. Journal of Bacteriology, 173(5):1817–1820, 1991.

[11] Rosario Lagunas. Sugar transport in saccharomyces cerevisiae. FEMS Microbiology Reviews, 1993.

[12] M J De Vit, J A Waddle, and M Johnston. Regulated nuclear translocation of the mig1 glucose repressor. Molecular Biology of the Cell, 8(8):1603–1618, 1997.

[13] S. Ozcan. Glucose sensing and signaling by two glucose receptors in the yeast saccharomyces cerevisiae. The EMBO Journal, 17(9):2566–2573, 1998.

[14] Jeong-Ho Kim and Mark Johnston. Two glucose-sensing pathways converge on rgt1 to regulate expression of glucose transporter genes in saccharomyces cerevisiae. Journal of Biological Chemistry, 281(36):26144–26149, 2006.

[15] Roman Wieczorke, Stefanie Krampe, Thomas Weierstall, Kerstin Freidel, Cornelis P Hollenberg, and Eckhard Boles. Concurrent knock-out of at least 20 transporter genes is required to block uptake of hexoses insaccharomyces cerevisiae. FEBS Letters, 464(3):123–128, 1999.

[16] Virve Vidgren, Laura Ruohonen, and John Londesborough. Characterization and functional analysis of the mal and mph loci for maltose utilization in some ale and lager yeast strains. Applied and Environmental Microbiology, 71(12):7846–7857, 2005.

[17] Hila Gingold, Disa Tehler, Nanna R. Christoffersen, Morten M. Nielsen, Fazila Asmar, Susanne M. Kooistra, Nicolaj S. Christophersen, Lise Lotte Christensen, Michael Borre, Karina D. Sørensen, Lars D. Andersen, Claus L. Andersen, Esther Hulleman, Tom Wurdinger, Elisabeth Ralfkiær, Kristian Helin, Kirsten Grønbæk, Torben Ørntoft, Sebastian M. Waszak, Orna Dahan, Jakob Skou Pedersen, Anders H. Lund, and Yitzhak Pilpel. A dual program for translation regulation in cellular proliferation and differentiation. Cell, 158(6):1281–1292, 2014.

[18] Luis Fernando Montaño-Gutierrez, Marc Sturrock, Iseabail Farquhar, Kevin Correia, Vahid Shahrezaei, and Peter S. Swain. A push-pull system of repressors matches levels of glucose transporters to extra-cellular glucose in budding yeast. bioRxiv, page 2021.04.20.440667, Apr 2021.

[19] S. Levy, M. Kafri, M. Carmi, and N. Barkai. The competitive advantage of a dual-transporter system. Science, 334(6061):1408–1412, 2011.

[20] Evert Bosdriesz, Meike T. Wortel, Jurgen R. Haanstra, Marijke J. Wagner, Pilar de la Torre Cortés, and Bas Teusink. Low affinity uniporter carrier proteins can increase net substrate uptake rate by reducing efflux. Scientific Reports, 8(1):5576, 2018.

[21] Frans Bianchi, Joury S. van’t Klooster, Stephanie J. Ruiz, and Bert Poolman. Regulation of amino acid transport in saccharomyces cerevisiae. Microbiology and Molecular Biology Reviews, 83(4):nil, 2019.

[22] Rob Patro, Geet Duggal, Michael I Love, Rafael A Irizarry, and Carl Kingsford. Salmon provides fast and bias-aware quantification of transcript expression. Nature Methods, 14(4):417–419, 2017.

[23] Michael I Love, Wolfgang Huber, and Simon Anders. Moderated estimation of fold change and dispersion for rna-seq data with deseq2. Genome Biology, 15(12):550, 2014.

[24] F. Pedregosa, G. Varoquaux, A. Gramfort, V. Michel, B. Thirion, O. Grisel, M. Blondel, P. Prettenhofer, R. Weiss, V. Dubourg, J. Vanderplas, A. Passos, D. Cournapeau, M. Brucher, M. Perrot, and E. Duchesnay. Scikit-learn: Machine learning in Python. Journal of Machine Learning Research, 12:2825–2830, 2011.

